# The postsynaptic MAGUK scaffold protein MPP2 organises a distinct interactome that incorporates GABA_A_ receptors at the periphery of excitatory synapses

**DOI:** 10.1101/2020.05.29.123034

**Authors:** Bettina Schmerl, Niclas Gimber, Benno Kuropka, Jakob Rentsch, Stella-Amrei Kunde, Helge Ewers, Christian Freund, Jan Schmoranzer, Nils Rademacher, Sarah A. Shoichet

**Affiliations:** Neuroscience Research Center, Charité – Universitätsmedizin Berlin, Germany; Advanced Medical BioImaging Core Facility – AMBIO, Charité – Universitätsmedizin Berlin, Germany; Institute of Chemistry and Biochemistry, Freie Universität Berlin, Germany; German Center for Neurodegenerative Diseases (DZNE), Berlin, Germany

## Abstract

Recent advances in imaging technology have highlighted that scaffold proteins and receptors are arranged in sub-synaptic nanodomains. The synaptic MAGUK scaffold protein MPP2 is a component of AMPA receptor-associated protein complexes and also binds to the synaptic cell adhesion molecule SynCAM1. Using super-resolution imaging, we now show that MPP2 and SynCAM1 are situated at the periphery of the postsynaptic density. In order to explore MPP2-associated protein complexes, we used a quantitative comparative mass spectrometry approach and identified multiple GABA_A_ receptor subunits among the novel synaptic MPP2 interactors. We further show that GABA_A_ receptors are found together with MPP2 in a subset of dendritic spines and thus highlight MPP2 as a scaffold molecule capable of acting as an adaptor molecule that links peripheral synaptic elements critical for inhibitory regulation to central structures at the PSD of glutamatergic synapses.

## Introduction

At postsynaptic sites on glutamatergic neurons, a complex arrangement of transmembrane receptors, scaffold molecules, and regulatory proteins enables the coordinated regulation of synaptic transmission (for reviews see [1-3]). Recent advances in imaging technology have highlighted that scaffold proteins and receptors are not distributed evenly throughout the postsynapse, but instead are arranged in sub-synaptic nanodomains [4]. These nanodomains are regions in which specific proteins are present at higher concentrations than in the surrounding areas, and they are individually regulated, functional units that are highly dynamic [5, 6]. The relevance of their regulation for synaptic function is becoming increasingly apparent (for reviews see [7-9]). Super-resolution imaging data from glutamatergic synapses suggest that such scaffold protein nanoclusters are responsible for concentrating glutamate receptors at particular sub-synaptic sites [5]. The incidence of these clusters seems to roughly scale with spine size, and clusters have been observed to undergo morphological plasticity [5, 10]. Sub-synaptic cluster dynamics at the postsynapse can also influence synaptic transmission, e.g. by affecting diffusion and/or trapping of neurotransmitter receptors [5, 6, 11].

We and others have recently demonstrated that MPP2 is a postsynaptic scaffold protein that is present in AMPA receptor-associated protein complexes [12-15]. Like PSD-95 and related molecules, MPP family proteins are members of the MAGUK family of scaffold molecules. Different from PSD-95 molecules, they do not bind directly to glutamate receptors or their auxiliary subunits. However, MPP2 binds directly to SynCAM1 synaptic cell adhesion molecules [13] that are positioned at the periphery of the PSD [16], suggesting that they may serve functions that differ from those of the core synaptic MAGUKs.

In this study, we have investigated this new postsynaptic MAGUK and its structural role at the PSD of glutamatergic synapses. We demonstrate that MPP2, like SynCAM1, sits at the periphery of the PSD and that it interacts with a unique set of proteins that differs significantly in composition from the set of PSD proteins binding to PSD-95 family synaptic MAGUKs. These novel interactions highlight the role of MPP2 in linking core synaptic components to transmembrane proteins and regulatory molecules with complementary functions at the periphery of the PSD. Importantly, we show that multiple GABA_A_ receptor subunits are among the proteins that bind preferentially to the MPP2 scaffold molecule, and we show that GABA_A_ receptors are found together with MPP2 and other classical PSD markers in a subset of dendritic spines. We thus highlight that the MPP2 scaffold molecule is capable of acting as an adaptor molecule that links important PSD protein complexes with elements critical for inhibitory regulation at the PSD of glutamatergic synapses.

## Results

### MPP2 and its interaction partner SynCAM1 are positioned at the periphery of the PSD

To investigate how MPP2 links the SynCAM1 cell adhesion molecules with the core PSD components, we first took advantage of diverse imaging strategies to comparatively analyse endogenous PSD-95, MPP2 and SynCAM1 proteins in primary rat hippocampal neurons (Fig 1a). Using dual-colour *d*STORM, we observe clusters of SynCAM1 in a ring-like arrangement of approximately 500 nm diameter (Fig 1b, upper panel) surrounding the PSD marked by PSD-95, which is in line with published data [17].

**Fig 1:**
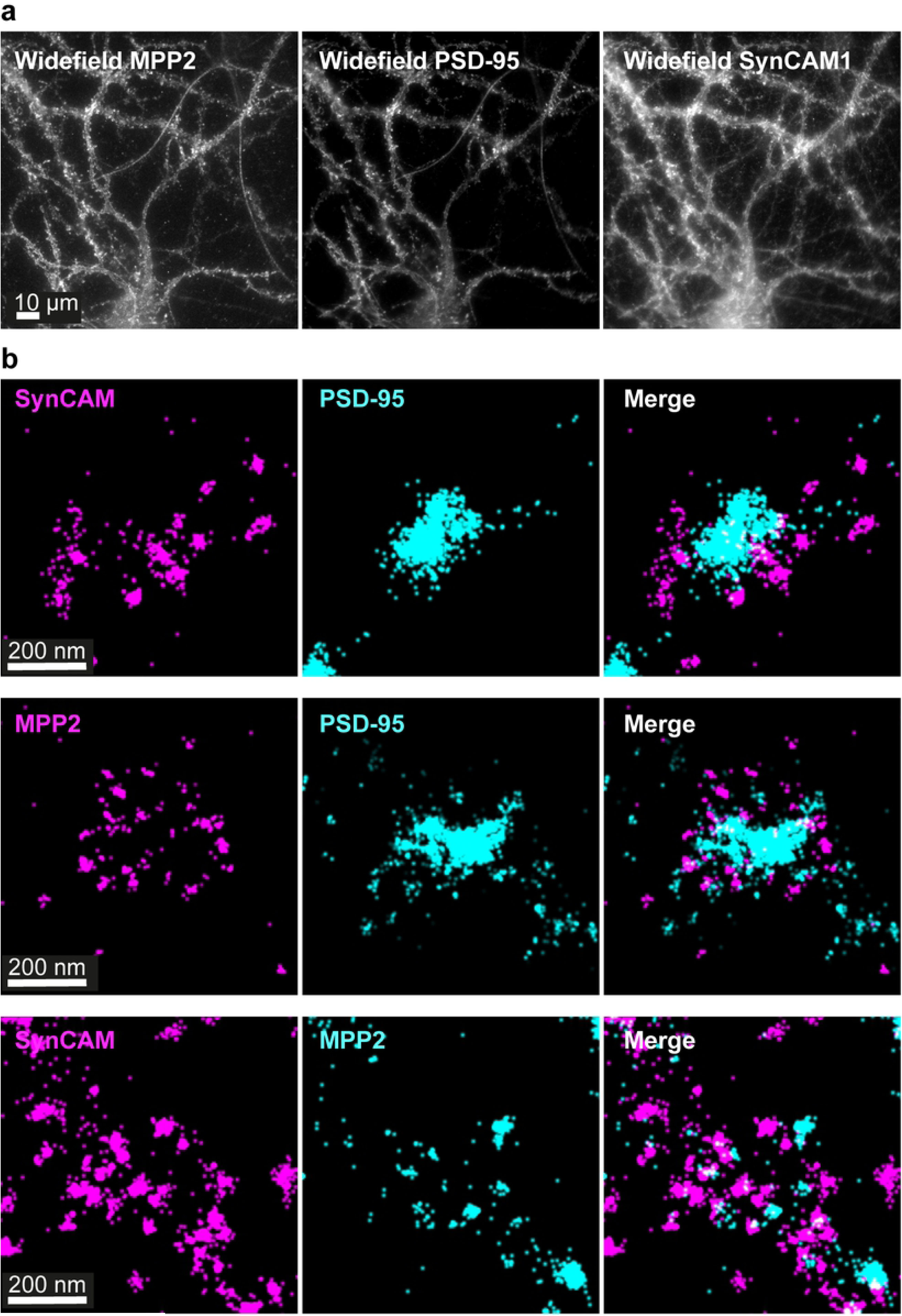
Clusters of SynCAM1 and MPP2 surround the postsynaptic density. (a) Widefield images of E18 primary rat hippocampal neurons fixed at DIV21 and stained for endogenous MPP2, PSD-95 and SynCAM1. (b) E18 rat hippocampal neurons were fixed at DIV21 and subjected to immunostaining for endogenous SynCAM1, PSD-95 and/or MPP2 proteins followed by dual-colour dSTORM imaging. Protein localisations were filtered according to Thompson accuracy [18], i.e. all localisations with accuracy below 20 nm were excluded. Top: SynCAM1 (magenta) clusters surround PSD-95 (cyan). Middle: Clusters of endogenous MPP2 (magenta) show a similar bracelet-like arrangement surrounding the PSD (PSD-95, cyan). Bottom: SynCAM1 (magenta) and MPP2 (cyan). For additional images please see supplements (S1 Fig, S2 Fig, S3 Fig).

Next, we examined the subcellular localisation of endogenous MPP2 and found a similar bracelet-like arrangement (approximately 500 nm in diameter), of small clusters of MPP2 surrounding postsynaptic densities as marked by PSD-95 (Fig 1b, middle panel). Further, when we stained for SynCAM1 in combination with MPP2, we observe that immunofluorescence of the two proteins is indeed arranged in a similar manner: at postsynaptic sites, we observe ring-like arrangements of SynCAM1 and MPP2 clusters that associate with each other and exhibit minor overlap (Fig 1b, lower panel).

### MPP2 is located in clusters at the periphery of the postsynaptic density

To assess whether this ring-like arrangement of MPP2 and SynCAM1 clusters surrounding PSD-95 is representative for the majority of synapses (and to avoid selection bias), a quantitative 3D super-resolution approach is necessary. We therefore tested whether we could resolve ring-like MPP2 and SynCAM1 structures also using 3D multicolour structured illumination microscopy (SIM). While offering less spatial resolution compared to *d*STORM, SIM inherently produces 3-dimensional data and can easily be adapted for more than two colour channels. Indeed, 3D SIM imaging is sufficient to observe similar planar bracelet-like cluster arrangements of SynCAM1 and MPP2 surrounding central PSD-95 labelled PSDs (Fig 2a).

**Fig 2:**
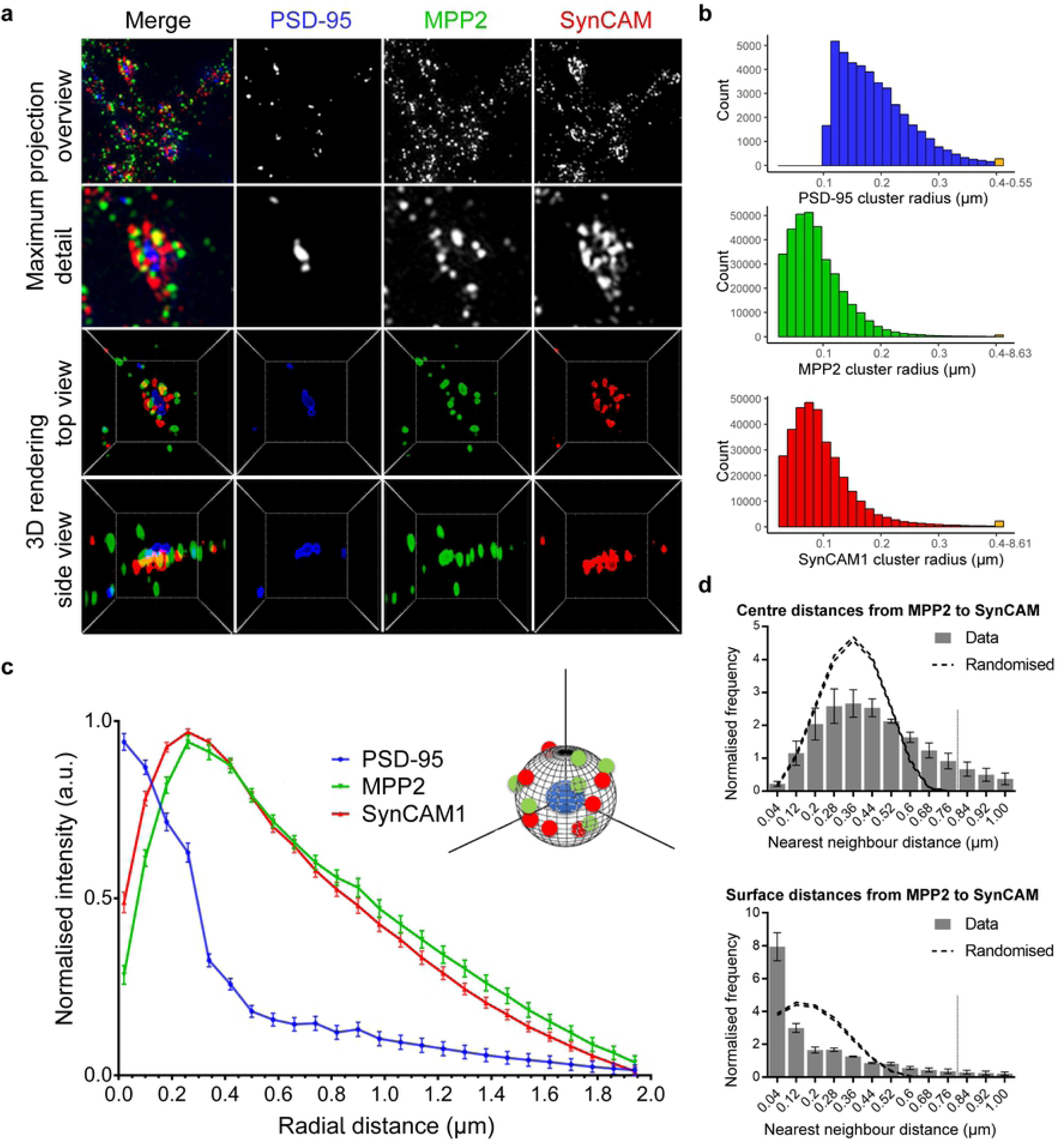
Clusters of MPP2 and SynCAM1 form bracelet-like arrangements at the edge of the postsynaptic density. Mature (DIV21) primary rat hippocampal neurons immunostained for endogenous PSD-95 (blue, second column), MPP2 (green, third column) and SynCAM1 (red, fourth column) and subjected to 3D structured illumination microscopy (3D SIM). (a) Most dendritic spines express all three proteins of interest (overview maximum projection, first row). A single synapse detail (second row) depicts the bracelet-like arrangements of MPP2 and SynCAM1 surrounding central PSD-95 puncta. A 3D rendering of that particular synapse in top (third row) and side view (fourth row) reveals that SynCAM1 and MPP2 clusters are arranged in an interlocked, bracelet-like form, surrounding a central cluster of PSD-95. Box sizes: overview = 7.7 µm, detail = 2.5 µm, 3D rendering = 2.8 µm. (b) Histograms illustrating the distribution of protein cluster sizes for PSD-95 (top, blue), MPP2 (middle, green) and SynCAM1 (bottom, red). Indicated radii were calculated based on extracted cluster volumes, assuming a spherical shape. The final bin in each histogram contains summarised data for cluster sizes greater than 400 nm. Histograms reflect clusters associated with ∼40,000 synapses (in 50 images from N = 3 independent experiments). (c) 3D radial intensity profiles of PSD-95, MPP2 and SynCAM1 signals in relation to the centres of PSD-95 clusters. Plot shows averaged normalised mean ± SEM from three independent experiments (∼40.000 synapses from 50 images). For details on the analysis, please see the methods. (d) Nearest neighbour (NN) analysis of MPP2 and SynCAM1 protein clusters after 3D segmentation. NN distances from MPP2 to the nearest SynCAM1 cluster were calculated from the cluster centres (upper panel, grey bars) and cluster surfaces (lower panel). Dashed lines represent the upper and lower envelopes of complete spatial randomness (CSR). CSR was calculated by randomly distributing MPP2 within the volume and SynCAM1 on the surface of spheres of 0.8 µm diameter as indicated by the grey dotted line (mean ± SEM, 95% confidence interval, 10 simulations per synapse, N = 3 independent experiments, ∼40.000 synapses from 50 images). See S4 Fig for NN analysis in the reverse direction.

Using semi-automated image segmentation (see methods) we quantitatively assessed the segmented object counts and radii as derived from the cluster volumes. In line with published data [6, 10] and our *d*STORM results, we find PSD-95 clusters over a range of expected sizes (Fig 2b, upper panel, blue). Interestingly, the majority of MPP2 (Fig 2b, middle panel, green) and SynCAM1 (Fig 2b, lower panel, red) clusters are smaller than ∼100 nm in radius.

Next, we quantitatively analysed the SynCAM1 and MPP2 protein distribution in relation to PSD-95 clusters, by measuring 3-dimensional radial distribution of all three proteins around the PSD centre defined by the PSD-95 signal. The 3D radial intensity profile of PSD-95 immunofluorescence intensity drops considerably at a radial distance of ∼250 nm (Fig 2c, blue curve), which is consistent with reported PSD sizes. The SynCAM1 signal (Fig 2c, red curve) is low at the centre of the postsynaptic density and highest towards the border of the PSD (radial distance of ∼250 nm), which is observable here in the steep decrease in PSD-95 signal. This data is in line with the idea that clusters of SynCAM1 define the edge of the PSD and the synaptic cleft [17]. Interestingly, the 3D radial intensity profile for MPP2 is almost identical to that of SynCAM1: we observe little fluorescence towards the centre of the postsynaptic density (radial distances below 250 nm) and the highest signal at the PSD border (Fig 2c, green curve). This quantitative result validates our qualitative super-resolution observation that MPP2, like SynCAM1, is distributed at the periphery of the PSD, around the core PSD protein PSD-95.

To assess the spatial relationship of the peripheral SynCAM1 and MPP2 protein clusters, we performed nearest neighbour (NN) analysis, which interrogates the nanoscale distances of the closest SynCAM1 cluster to each MPP2 cluster, and *vice versa*. This NN analysis was performed by assessing both the distances from centre-to-centre and from surface-to-surface for each 3-dimensional object. The distance distributions were quantitatively compared to a simulated random distribution within the known volume of a postsynapse [19]. The centre-to-centre NN analysis showed that most centres of MPP2 clusters have an NN distance of 200 nm - 500 nm to centres of SynCAM1 clusters (Fig 2d, upper panel). Similar results were obtained when analysing the centre-to-centre distances of SynCAM1 clusters to the nearest MPP2 cluster (see S4 Fig), showing that the two proteins do not form one uniform cluster. The obtained range of NN distances rather corresponds to the sum of both cluster radii (average cluster sizes are below 200 nm; Fig 2b, middle and lower panel), suggesting a juxtapose association of the proteins.

To test whether the surfaces of MPP2 and SynCAM1 clusters overlap with each other, we performed a surface-to-surface NN analysis, which showed a significant accumulation of MPP2 clusters at very small distances to SynCAM1 clusters and *vice versa* (Fig 2d, lower panel). This accumulation is very prominent regarding the MPP2 clusters that are located around SynCAM1 clusters (Fig 2d, lower panel), however, it is less prominent when analysed inversely (see S4 Fig). This indicates that most MPP2 clusters are associated with SynCAM1 clusters but not *vice versa*, suggesting the existence of an additional SynCAM1 pool that is independent from the postsynaptic MPP2, which is in line with the fact that SynCAM1 is also present at presynaptic sites. In summary, these data show that MPP2 and SynCAM1 clusters are not spatially identical but tightly juxtaposed at the periphery of the PSD.

Although clusters of SynCAM1 and MPP2 are not spatially identical, their close association offers sufficient chances for molecular interaction with each other. At the same time, it highlights the possibility that other surfaces remain accessible for interactions with other proteins. This is consistent with the idea that the MAGUK scaffold protein MPP2, like PSD-95, serves as a protein-protein interaction hub and that it is able to mediate the indirect connection of multiple proteins through diverse, domain-specific protein-protein interactions. In summary, our super-resolution data provide evidence that MPP2 and SynCAM1 are postsynaptic proteins in a close sub-synaptic arrangement sitting at the periphery of postsynaptic densities, rather than being central components of the PSD, and thus highlight the potential for the scaffold protein MPP2 to mediate formation of complexes that are distinct from the central nanodomains orchestrated by the neighbouring MAGUK PSD-95.

### The C-terminal SH3GK domains of MPP2 and PSD-95 interact with distinct synaptic proteins

The observed peripheral synaptic localisation of MPP2 relates to that of its PDZ ligand binding partner SynCAM1 [13, 16]. PSD-95 is located in central sub-synaptic nanodomains that likewise correlate with the localisation of its PDZ domain-binding partners, i.e. glutamate receptors and auxiliary proteins [20-23]. The SH3 and GK domains, which are typically located at the C-terminus of MAGUK scaffold proteins (for overview of MPP2 and PSD-95 domain architecture see Fig 3a), also participate in scaffold complex formation. Importantly, it has been shown that the GK domain of MAGUK proteins is an inactive guanylate kinase [24] that has evolved into an important protein interaction domain. An intramolecular interaction between the SH3 and GK domains of MAGUKs has been well-characterised, and several studies also support the idea that these domains of the MAGUK protein PSD-95 are involved in regulated multi-protein complex formation [25, 26]. While numerous binding partners for this region of PSD-95 have been described, interactors specific for the SH3GK domain of MPP2 have not been identified so far.

**Fig 3:**
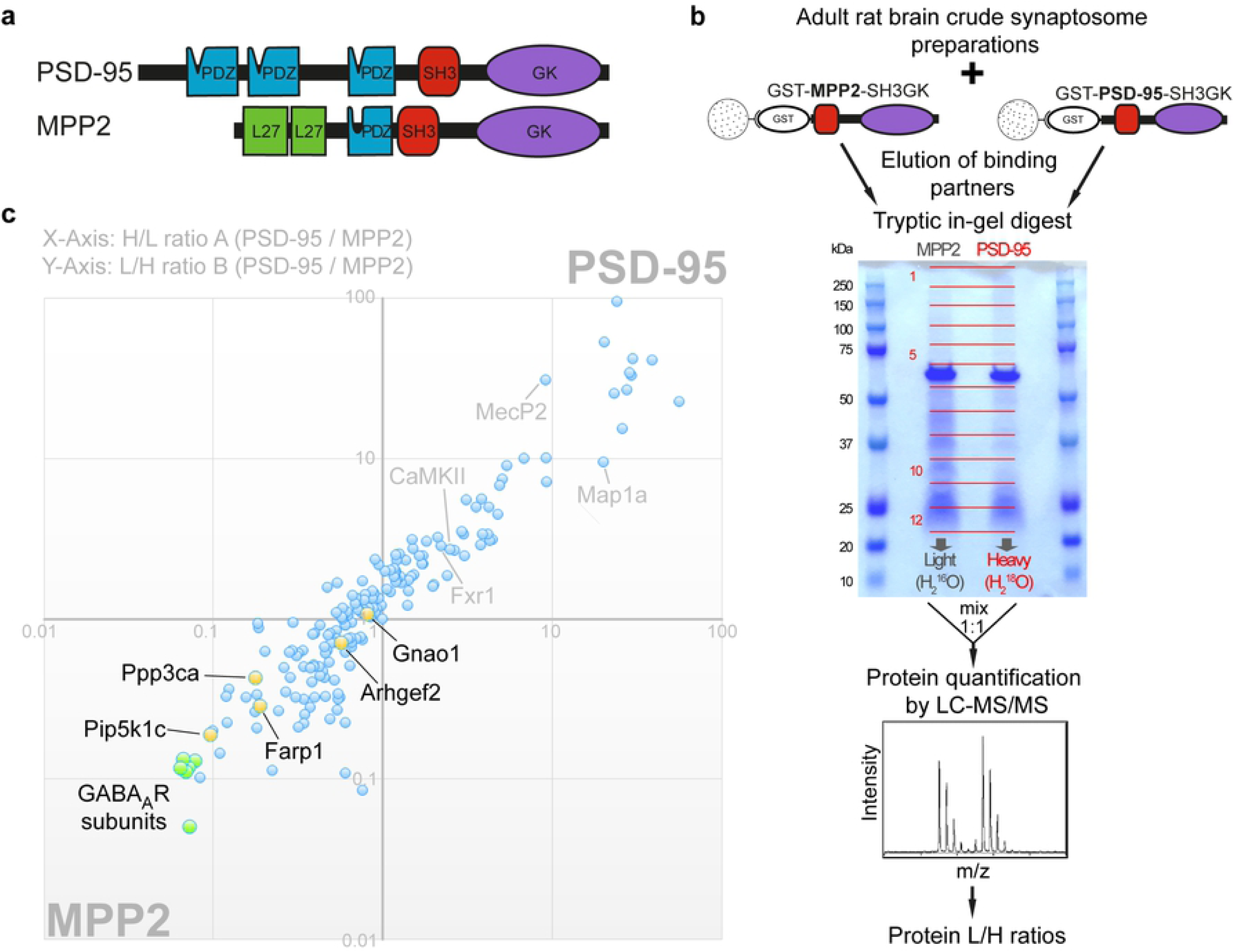
Identification of interactors that differentially bind to the C-terminal SH3GK modules of MPP2 and/or PSD-95. (a) Schematic domain structures of PSD-95 and MPP2 to scale and aligned by their central PDZ domain. Both proteins contain two N-terminal domains (PDZ1+PDZ2 for PSD-95 and two L27 domains for MPP2) in addition to the C-terminal ‘MAGUK core’ domains PDZ-SH3-GK. Note the differences in the length of the ‘linker’ between PDZ and SH3 domain and of the ‘hook’ between SH3 and GK domains. (b) Schematic representation of the quantitative LC-MS/MS experiment using ^16^O/^18^O-labelling to identify differential interactors from adult rat brain crude synaptosomal preparations by GST pull-down of bacterially expressed GST-MPP2-SH3GK or GST-PSD-95-SH3GK. (c) GST pull-downs were performed in duplicates with inverted labelling and 188 interacting proteins were identified and quantified by mass spectrometry passing our threshold settings. PSD-95 / MPP2 protein ratios from both replicates A and B (normalised by the ratio of GST) are plotted against each other. Proteins in the first quadrant indicate preferential enrichment to PSD-95 (ratios >> 1), while proteins in the third quadrant indicate preferential enrichment to MPP2 (ratios << 1). Proteins with ratios of ∼1 show no preferential binding, and thus reflect equal binding to both baits or background proteins that were not fully removed by the washing steps. Selected novel potential interaction partners were validated by co-IP (yellow, see also S5 Fig). The most significantly enriched proteins to the GST-SH3GK construct of MPP2 (green cluster) are seven different GABA_A_ receptor subunits.

Given the observed peripheral localisation of MPP2, we hypothesised that identification of its C-terminal interaction partners might illuminate protein complexes that differ from those organised by the central PSD-95 scaffold molecule. To explore this idea, we performed a comparative and quantitative mass spectrometry analysis (see Fig 3b for experimental design). Using bacterially expressed GST-MPP2-SH3GK and GST-PSD-95-SH3GK, we pulled down SH3GK-binding proteins from adult rat crude synaptosome preparations. Interacting proteins were eluted from the beads and separated by SDS-PAGE. Enzymatic ^16^O/^18^O-labelling was used for relative quantification of proteins by nanoLC-MS/MS analysis. In replicate A, proteins enriched by MPP2-SH3GK carried the naturally highly abundant ^16^O isotope, while proteins enriched by PSD-95-SH3GK were labelled by ^18^O using H_2_^18^O during tryptic in-gel digestion (see Fig 3b). In replicate B, labels were switched. In total, we reproducibly identified and quantified 188 proteins (see Source Data). Plotting the protein heavy/light ratios (H/L) from replicate A (Fig 3c, X-axis) against the light/heavy ratios (L/H) from replicate B (Fig 3c, Y-axis) shows proteins enriched to PSD-95-SH3GK in the first quadrant and proteins enriched to MPP2-SH3GK in the third quadrant (Fig 3c). Background proteins (not completely washed from the beads) and proteins which bind to both SH3GK constructs show ratios of about 1. From all identified proteins, 83% have been previously reported in human and/or mouse postsynaptic density preparations [27], confirming the validity of our approach for identifying true PSD proteins. Further evidence illustrating the strength of our strategy is the fact that several known interactors of PSD-95 (including e.g. Map1a, MecP2, CaMKII and Fxr1) were identified among proteins enriched in the GST-PSD-95-SH3GK pull-down (Fig 3c).

Importantly, proteins found consistently in the MPP2 pull-downs reflect putative novel synaptic MPP2 binding proteins. Of the newly identified proteins present in the GST-MPP2-SH3GK pull-down samples (see S1 Table Source Data), most have been found in PSD preparations before [27], which is in line with our previous work highlighting MPP2 as an important postsynaptic scaffold.

Several putative novel MPP2 interaction partners were selected for validation and further study (see Fig 3c, highlighted proteins; see also Table 1 for more detail). We demonstrated that both Gnao1 and Arhgef2, proteins involved in signalling cascades relevant for postsynaptic function [28, 29], interact with MPP2 in co-immunoprecipitation assays (see S5 Fig). We also validated a clear interaction between MPP2 and the membrane-associated synaptic proteins Farp1 and Pip5k1c (S5 Fig).

**Table 1:** Validated novel interaction partners for MPP2 (for co-immunoprecipitation data see Fig 4a and S5 Fig)

| Name | Uniprot Accession ID | Description | Remarks |
| --- | --- | --- | --- |
| Gnao1 | GNAO_RAT | Guanine nucleotide-binding protein G(o) subunit alpha | transducer in transmembrane signalling systems |
| Arhgef2 | ARHG2_RAT | Rho guanine nucleotide exchange factor 2 | Interacts with AMPA receptors |
| Farp1 | FARP1_RAT | FERM, ARHGEF and pleckstrin domain-containing protein 1 | SynCAM interactor |
| Pip5k1c | PI51C_RAT | Phosphatidylinositol 4-phosphate 5-kinase type-1 gamma | Binds to FERM domains, activated by Rho / Rho GEF signaling |
| Ppp3ca | PP2BA_RAT | Serine/threonine-protein phosphatase 2B catalytic subunit alpha isoform | Calcineurin subunit; known GABA<sub>A</sub> R interactor |
| GABAA α1 | GBRA1_RAT | Gamma-aminobutyric acid receptor subunit alpha-1 | Predominant GABA<sub>A</sub> subunit in the brain |

Importantly, in addition to revealing previously unknown binding partners for MPP2, our comparative quantitative MS strategy provided us with important information on how the MPP2 interactome relates to that of PSD-95. Of particular interest, the set of proteins most significantly enriched in the GST-MPP2-SH3GK pull-down comprised multiple gamma-aminobutyric acid (GABA) A receptor subunits (highlighted in green in Fig 3c; see also Table 1 and Source Data). Seven different GABA_A_ subunits, namely α1, α2, α4, β1, β2, β3, and δ, were enriched more than seven-fold. In combination, these subunits are able to form complete hetero-pentameric receptors [30, 31], illustrating the potential for MPP2 molecules to interact at multiple sites with a fully functional GABA_A_ receptor. These results are striking, as GABA_A_ receptors are not known to be expressed at high levels at the PSD of glutamatergic synapses.

We validated the interaction between MPP2 and the GABA_A_ α1 subunit by co-immunoprecipitation (see Fig 4a and Table 1). In this context, it is also interesting that MPP2 also interacts with the calcium-dependent, calmodulin-stimulated protein phosphatase (Calcineurin) subunit Ppp3ca (see Fig 3c, Table 1, and S5 Fig), which is known to influence GABA_A_ receptor signalling [32]. Together these data support the novel idea that MPP2 could be involved in GABA_A_ receptor-mediated processes at the PSD of glutamatergic synapses.

**Fig 4:**
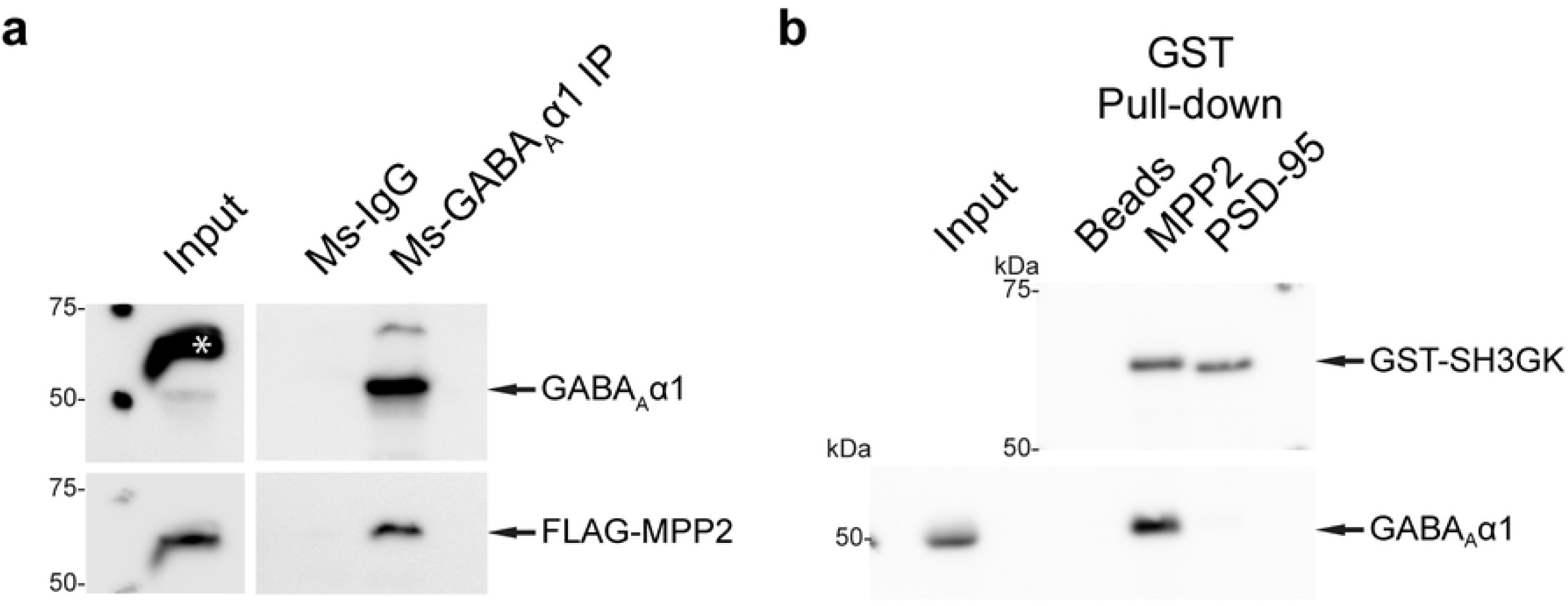
Validation of GABA_A_ α1 as novel interaction partner of MPP2. (a) Untagged GABA_A_ α1 was overexpressed in CHL V79 cells together with FLAG-tagged MPP2 and subjected to pull-down with αGABA_A_ α1 or normal Ms IgG. Co-purification of FLAG-MPP2 was detected by Western blot with αFLAG and αGABA_A_ α1 primary and α-native-Ms-IgG (Veriblot) secondary antibodies. Asterisk (*) marks re-emergence of αFLAG-HRP signal from the previous exposure. (b) Bacterially expressed GST-MPP2-SH3GK and GST-PSD-95-SH3GK were incubated with crude brain synaptosome preparations. After GST pull-down, compared to bead control, GABA_A_ α1 was efficiently enriched in the GST-MPP2-SH3GK pull-down, as detected by western blot with αGABA_A_ α1 and αGST antibodies.

### GABA_A_ receptor subunits are novel synaptic interaction partners of MPP2

The proteins most significantly enriched in the MPP2-SH3GK pull-down were multiple GABA_A_ receptor subunits. To further investigate this unexpected result, we overexpressed GABA_A_ α1 together with full-length FLAG-MPP2. Upon pull-down of the receptor subunits with Ms-αGABA_A_ α1 antibody, we could detect MPP2 in the precipitate (Fig 4a), suggesting that GABA_A_ α1 is indeed a new binding partner for MPP2. Moreover, our comparative pull-down experiments using GST-tagged SH3GK domains of MPP2 and PSD-95 validate our mass spectrometry data (Fig 3) and demonstrate that MPP2 – in contrast to PSD-95 – binds effectively to GABA_A_ α1: while the MPP2 SH3-GK domain efficiently pulls out the endogenous GABA_A_ α1 from crude synaptosome preparations, the PSD-95 SH3-GK domain does not (Fig 4b).

### MPP2 co-localises with endogenous GABA_A_ α1 in a sub-population of dendritic spines

Our *in vitro* experiments clearly show that GABA_A_ receptor α1 subunits preferentially interact with the C-terminal SH3GK module of MPP2. In light of the fact that GABA_A_ receptors are best known for their importance at inhibitory synapses where Gephyrin is the predominant scaffold (for review see [33]), this result was surprising. To explore the idea that GABA_A_ receptors interact with MPP2 at glutamatergic synapses, we immunostained dissociated rat primary hippocampal neurons for the endogenous proteins. Neuronal cultures were fixed at DIV21 and stained for GABA_A_ α1 and MPP2 together with the dendritic marker MAP2 (microtubule-associated protein 2) and the postsynaptic density marker Homer1, with respective primary and secondary antibodies. MPP2 (Fig 5, cyan) is present in almost all dendritic spines positive for the PSD marker Homer1 (Fig 5, yellow). Upon analysis of secondary dendrites (Fig 5b), we observed GABA_A_ α1 signals (Fig 5, magenta) not only along MAP2-positive dendritic branches, where expected (Fig 5, blue), but also co-localising directly with MPP2 and Homer1 in a subset of dendritic spines (Fig 5b and c). More detailed examination of selected spines via line plot analysis of fluorescence intensity (Fig 5d) confirmed three different patterns of protein expression among the spines in our cultures: synapses positive for all three proteins of interest (Fig 5d i), excitatory synapses positive for MPP2 and Homer1 (Fig 5d ii), and inhibitory synapses at the dendrite expressing GABA_A_ α1 but negative for MPP2 and Homer1 (Fig 5d iii).

**Fig 5:**
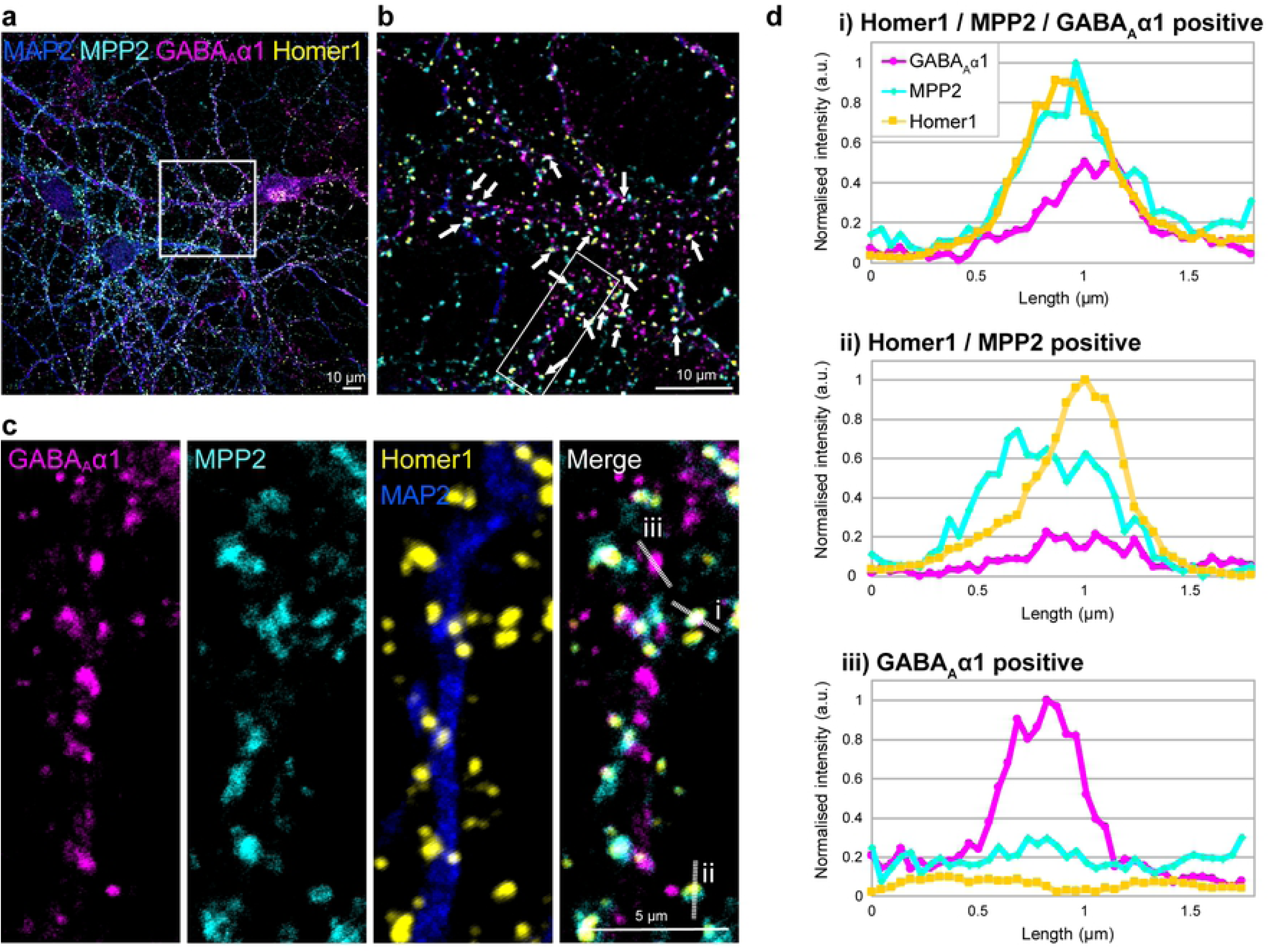
GABA_A_ receptor subunits α1 co-localise with MPP2 in a subset of dendritic spines. (a) Primary E18 rat hippocampal neurons were fixed at DIV21 and immunostained for endogenous proteins MAP2 (microtubule-associated protein 2), MPP2, GABA_A_ α1 and Homer1 using respective primary and Alexa fluorophore-coupled secondary antibodies, and visualised by confocal microscopy. Box in this single-plane overview indicates location of detail image in b. (b) Maximum projection composite of four-colour confocal immunofluorescence images acquired with 5x zoom depicts primary and secondary dendrite branches of a mature (DIV21) hippocampal neuron. Arrows indicate several dendritic spines where GABA_A_ α1 co-localises with Homer1 and MPP2. Box indicates further zoom in c. (c) Detail zoom of a secondary dendrite branch. Markers at positions a, b and c indicate line plot measurement locations shown in d. (d) Normalised fluorescence intensity line plot quantification of exemplary dendritic spines as shown in c. Along the dendrite we find synapses in which GABA_A_ α1, Homer1 and MPP2 clearly co-localise (i) and synapses positive for Homer1 and MPP2 with no GABA_A_ α1 immunofluorescence (ii), as well as solely GABA_A_ α1 positive punctae, likely representing inhibitory synapses at the dendrite (iii).

Together with our *in vitro* data, these results suggest that MPP2 and GABA_A_ receptors interact in a sub-population of excitatory synapses and that MPP2 acts as a scaffold protein potentially involved in the recruitment and anchoring of GABA_A_ receptors at these synapses.

## Discussion

We are interested in how sub-synaptic nanodomains at glutamatergic synapses are organised such that they can orchestrate synaptic function, which is highly dynamic. Here we show that PSD-95 and MPP2, two related synaptic MAGUKs, are close, but distinctly localised at the PSD of glutamatergic synapses. Using super-resolution imaging strategies, we observe that SynCAM1 and MPP2 are located in juxtapose association towards the outside of the PSD, with both proteins surrounding central PSD-95 protein complexes at radial distances that reflect a peripheral PSD localisation, given typical expected PSD sizes [19].

These observations, combined with the fact that MPP2 interacts directly with the peripheral SynCAM1 (but not the more central AMPAR-auxiliary subunits, TARPs), led us to pursue the idea that MPP2 may act as a scaffold for protein complexes that are distinct from those at the core of the PSD. Using a comparative mass spectrometry approach, we demonstrate here that the SH3GK domains of MPP2 and PSD-95, which are structurally similar, indeed interact with distinct sets of cytosolic and membrane proteins present within dendritic spines. Importantly, we identified several novel MPP2-interacting proteins, including multiple GABA_A_ receptor subunits as well as signalling molecules with established roles in the regulation of inhibitory transmission. Moreover, in hippocampal neuronal cultures, we observe a subset of glutamatergic synapses that clearly express both GABA_A_ receptors and MPP2. Together, these data indicate a role for MPP2 as a scaffold protein that organises sub-synaptic nanodomains distinct from those defined by PSD-95, and highlight its potential to act as a mediator of inhibitory signalling at glutamatergic synapses.

Our combined imaging strategies illustrate that MPP2 and SynCAM1 sit directly next to each other towards the outside of the PSD, positioned optimally to regulate the formation of peripheral sub-synaptic nanodomains, and our comparative quantitative mass spectrometry data provide further information on the nature of these protein complexes. Importantly, well-known PSD-95-interacting proteins, including e.g. Map1a and MecP2, were highly enriched in the PSD-95-SH3GK pull-downs, whereas Farp1, a well-characterised SynCAM1 interactor and modulator of SynCAM-mediated processes [34], was found to be significantly enriched in our MPP2-SH3GK pull-downs. These data in particular illustrate the utility of our strategy for identifying unknown MPP2 interactors of potential importance.

Among the crude synaptosome proteins present in the MPP2-SH3GK pull-down, we consistently identified seven GABA_A_ receptor subunits, i.e. more than one-third of all known subunits[35]. The fact that so many GABA_A_ receptor subunits were in the top hits among MPP2-associated peptides supports the idea that the MAGUK-GABA_A_ receptor association is MPP-specific and that these receptors do not generally bind with high affinity to the SH3GK domains of other synaptic MAGUKs. In co-immunoprecipitation experiments we could confirm a direct interaction of GABA_A_ α1 with MPP2. Moreover, in our primary hippocampal neuron cultures, we also detect endogenous GABA_A_ receptors in a subset of Homer1-positive glutamatergic synapses that express MPP2, providing support for the idea that there may be a functional role of MPP2-GABA_A_ receptor protein complexes in dendritic spines.

GABA_A_ receptors mediate inhibitory transmission onto dendrites at diverse locations of the dendritic arbour (for reviews see [32, 33, 36]). Detailed studies on their cellular localisation have revealed that GABA_A_ receptor complexes are present not only on dendritic shafts but also in dendritic spines and in close proximity to PSDs of glutamatergic synapses [37]. Using electron microscopy, this perisynaptic localisation of GABA_A_ receptor subunits has also been monitored by other groups in a developmental context [38], and more recently, the importance of GABA_A_ receptors within dendritic spines and their role in regulating Ca^2+^ influx [39] and general excitatory signal transmission [40] has become a topic of considerable interest (for review see [36]). Recent studies also highlight that GABA_A_ receptor mobility between inhibitory synapses is an important mechanism by which inhibitory synaptic currents are regulated, and that this process can be modulated by activation of ionotropic glutamate receptors and subsequent trapping of desensitized GABA_A_ receptors at glutamatergic synapses [41]. Clearly, there are multiple functional links between GABA_A_ receptor signals and glutamatergic transmission at the PSD. However, the physical interactions that enable these connections have not been investigated to date. Our imaging studies highlight that MPP2 is positioned optimally – next to the core components of glutamate receptor signalling complexes, but at the periphery of the PSD – to play a structural role in orchestrating the complex formation that would be required to enable these dynamic links between glutamatergic transmission and GABA_A_ receptor signalling.

We have previously demonstrated that MPP2 is linked to AMPA receptor protein complexes via SH3GK domain-mediated interactions with the core AMPA receptor-associated scaffold proteins PSD-95 and GKAP. Here we show that the same domains are responsible for specific interactions with GABA_A_ receptors. Interestingly, the L27 domains at the N-terminus of MPP2 molecules are well known for their role in mediating MPP2 dimerisation [42]. In this context, our data, which demonstrates a physical link between GABA_A_ receptors and MPP2 at glutamatergic synapses, lead us to propose a model in which MPP2 multi-molecular complexes serve as adaptors that enable crosstalk between GABA_A_ receptor-mediated inhibitory regulation and glutamatergic transmission in dendritic spines (see Fig 6). Our model is further supported by the fact that one of the other novel MPP2 interaction partners identified in this study has previously been associated with inhibitory signalling through GABA_A_ receptors: the calcium- and calmodulin-dependent serine/threonine protein phosphatase (Calcineurin) subunit Ppp3ca directly interacts with GABA_A_ receptors [43-45].

**Fig 6:**
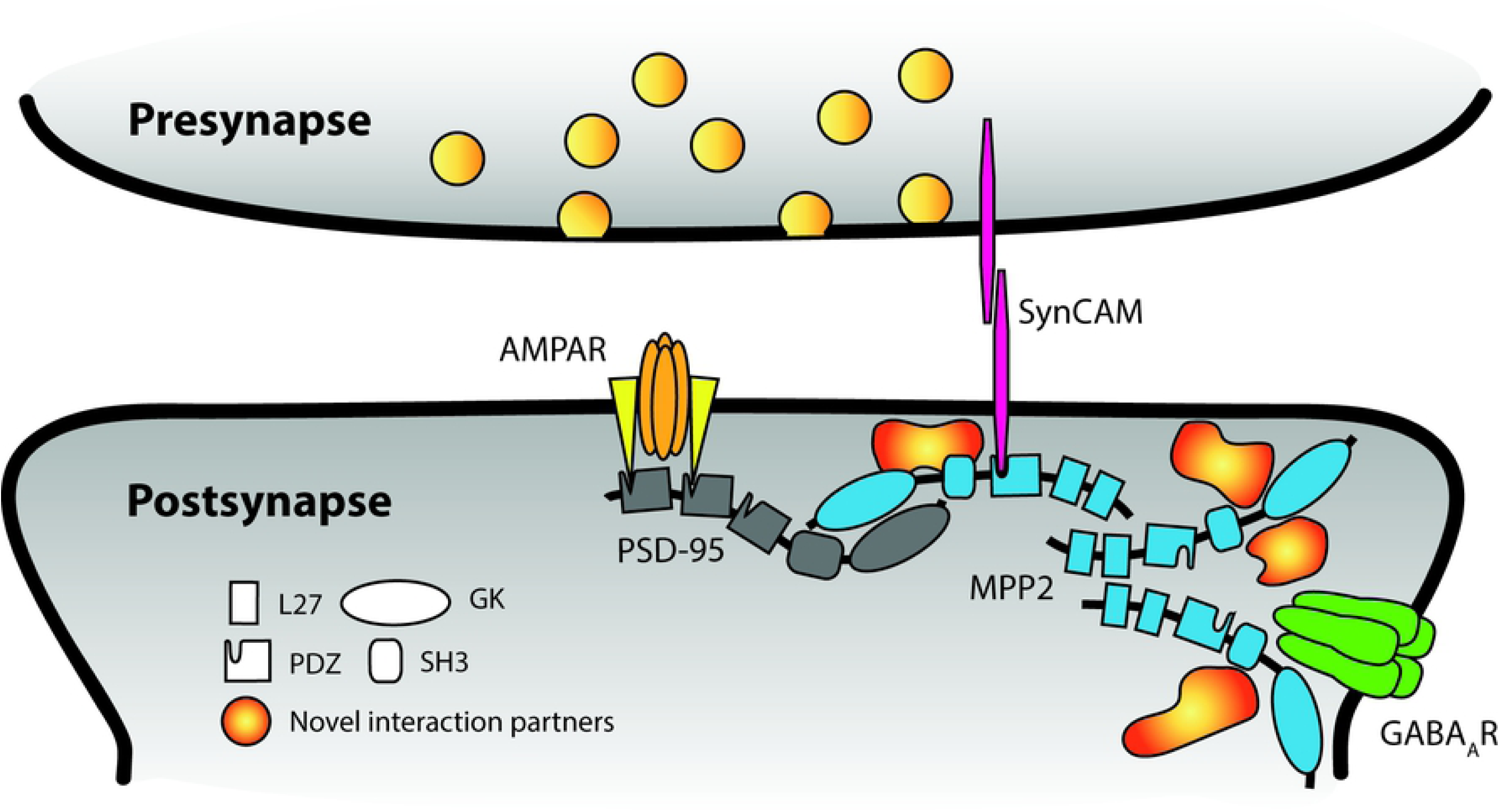
Schematic summary of novel MPP2 interactors. Our data indicate an important role of MPP2 (blue) in the sub-synaptic compartmentalisation of dendritic spines by connecting central components of AMPA receptor (orange) complexes, like TARPs (yellow) and PSD-95 (grey) not only to cell adhesion proteins like SynCAM (magenta) and other scaffold and regulatory proteins (like the novel interaction partners PIP5k1c or Ppp3ca), but most importantly inhibitory GABA_A_ receptors (green). For MAGUK proteins the individual domain structure is indicated.

In summary, our study provides insights into the physical interactions that mediate complex formation in dendritic spines. We demonstrate that the MPP2 scaffold protein serves to link core proteins of glutamatergic synapses with GABA_A_ receptors and associated signalling molecules in dendritic spines, and thereby illuminate its potential to facilitate crosstalk between excitatory and inhibitory transmission at the PSD of glutamatergic synapses.

## Materials and Methods

### Primary neuronal cultures

Primary hippocampal neurons were prepared as described before [13] with protocols approved by ‘Landesamt für Gesundheit und Soziales’ (LaGeSo; Regional Office for Health and Social Affairs; permit number T0280/10) in Berlin. Briefly, E18 Wistar rat pups were decapitated, hippocampi isolated and digested with Trypsin/EDTA (Lonza). Digest was stopped with DMEM/10 % FBS (Biochrom), followed by washing in DMEM (Lonza). Tissue was then dissociated and plated at ∼10^5^ cells per cm^2^ in neuron culture medium (Neurobasal (Lifetech) supplemented with B27 (Gibco) and 500 µM L-glutamine (Lonza)) onto coverslips coated with poly-D-Lysine (0,2 mg/ml, Sigma) and Laminin (2 µg/ml, Sigma) and maintained in a humidified incubator (37°C, 5% CO_2_).

### Immunocytochemistry/Immunofluorescence

Primary rat hippocampal neurons were fixed at DIV21 with 4 % PFA/PBS for 10 min at RT, washed thrice for 10 min with PBS, followed by 45 min quenching at RT with 50 mM NH_4_Cl to reduce auto-fluorescence. After washing with PBS, cells were permeabilised with 0.2 % Triton-X/PBS for 5 min and washed with PBS. For dSTORM microscopy cells were additionally treated with Image-IT Signal Enhancer (Thermo Fisher) for 30 min at RT and three washes with PBS. Cells were then blocked for at least 1 hr at RT with blocking solution (4% BSA/PBST). Primary antibodies were diluted 1:250 in blocking solution (1:500 for confocal microscopy) and incubated over night at 4°C, followed by incubation with desired secondary antibodies at 1:1000 dilution in blocking solution for 1 hr at RT and for *d*STORM custom-labelled secondary antibodies at 1 µg/ml (∼7 nM) in blocking solution for 20 min, followed by post-fixation and quenching as above. After final washing with PBS, coverslips were mounted with Fluoromount G (SBA) or Vectashield (H1000, Vector Laboratories) for confocal and SIM imaging, respectively.

#### Primary antibodies

αMPP2 (rabbit, ab97290, Abcam), αMAP2 (guinea pig, 188004, Synaptic Systems), αPSD-95 (mouse, 75-028, NeuroMab), αvGlut1 (guinea pig, 135 304, Synaptic Systems), αGABA_A_ α1 (mouse, 75-136, NeuroMab), αSynCAM (chicken, CM004-3, MBL), GFP-Booster (nanobody, gba488-10, Chromotek)

#### Secondary antibodies

αMouse Alexa Fluor 405 (A-31553, Invitrogen), αGuinea pig Alexa Fluor 405 (ab175678, Abcam), αGuinea pig DyLight405 (706-475-148, Dainova), αRabbit Alexa Fluor 488 (A-21441, Invitrogen), αChicken Alexa Fluor 488 (703-545-155, Jackson Immuno Research), αMouse Alexa Fluor 568 (A-11031, Life Technologies), αRabbit Alexa Fluor 568 (A-11036, Invitrogen), αRabbit (111-005-003, AffiniPure), αMouse (715-005-151, AffiniPure), αMouse Alexa Fluor 647 (715-605-150, Jackson Immuno Research), αChicken Alexa Fluor 647 (103-605-155, Dianova).

### Confocal Microscopy

Cells were fixed and stained as described above and imaged with a Leica TCS SP5 II confocal laser scanning microscope run with LAS AF X scan software (Leica Microsystems, Wetzlar, Germany). Image stacks were acquired with a 63x oil immersion objective at 2x zoom for overviews and 5x zoom for details at 2048×2048 px as 5-7 planes with 0.4 µm step size in Z. Image analysis was performed in Fiji/ImageJ[46, 47]. The three highest intensity planes for Homer1 were selected and maximum projected. Then, line plot profile measurements were taken for individual spines for every channel. Data was then 0/1 normalised across all acquired values within one channel.

### Dual-colour *d*STORM imaging

#### Labelling of antibodies

Secondary antibodies (goat αRabbit, 111-005-003, AffiniPure or donkey αMouse, 715-005-151, AffiniPure) were diluted 1:10 in labelling buffer (0.2 M NaHCO_3_, pH 8.3). Cy3b NHS Ester (PA63101, Life Sciences) was added to the diluted antibodies in 10-fold molar excess. The samples were incubated for 1 hr at room temperature. To stop the reaction, 100 mM Tris pH 8.0 was added. Zebra spin desalting columns (8989, Thermofisher) were equilibrated with PBS. The samples were added to the column and centrifuged at 1000 *g* for 2 min. The filtrate was added to a second column and centrifuged at 1000 *g* for 2 min.

#### Image acquisition

All samples were imaged using a Vutara 352 super-resolution microscope (Bruker) equipped with a Hamamatsu ORCA Flash4.0 sCMOS camera for super-resolution imaging and a 60x oil immersion TIRF objective with a numerical aperture of 1.49 (Olympus). Immersion Oil 23°C (#1261, Cargille) was used. Samples were mounted onto the microscope in GLOX buffer (1.5% β-mercaptoethanol, 0.5% (v/w) glucose, 0.25 mg/ml glucose oxidase and 20 µg/ml catalase, 150 mM Tris-HCl pH 8.8), illumination at a laser-power density of 5.5 kW/cm2 using a 637 nm laser for Alexa Fluor 647 or a 561 nm laser at a laser-power density of 4.6 kW/cm^2^ for Cy3b. Images were collected with 20 ms acquisition time. Per probe (Cy3b or Alexa Fluor 647) 10,000 frames were acquired. Acquired raw data were localized using SRX (Bruker). Localisations were filtered according to Thompsom accuracy [18], i.e. all localisations with accuracy worse than 20 nm were excluded. Localisations were rendered as Gaussian distributions with a constant width of 20 nm. Alignment of colour channels and drift correction were performed in SRX using Tetraspeck beads (Thermofisher, T7279). Supplemental Figures 1-3 were prepared using the ScientiFig Plugin for Fiji/ImageJ [48].

### Structured Illumination Microscopy

#### Sample preparation

Primary hippocampal rat neurons (DIV21) stained for the presynaptic marker vGlut1 and the postsynaptic proteins PSD-95, SynCAM1 and MPP2 as described above.

#### Image acquisition

Targets were selected based on the SynCAM1 signal. Three-dimensional SIM images were acquired with the OMX V4 Blaze system (GE Healthcare), using the 405 nm, 488 nm, 568 nm and 647 nm laser lines, a 60x 1.42 N.A. oil objective (Olympus), an immersion oil with a refractive index of 1.518 and standard emission filters at 125 nm z-sectioning. Multi-colour registration with an error below 40 nm was done using 100 nm fluorescent beads (TetraSpeck, T7284, Thermo Fisher Scientific). Images were acquired with the DeltaVision OMX acquisition software (GE Healthcare) and images were reconstructed with softWoRx (GE Healthcare).

### SIM Image Analysis

#### Segmentation

Image segmentation was performed in Arivis Vision 4D (Arivis AG, Munich, Germany). MPP2, SynCAM1 and vGlut1 clusters were identified with histogram-based threshold procedures (Otsu’s and Yen’s method). PSD 95 clusters and their centres were identified with the built in “Blob Finder” tool, a combination of automatic seed finding and watershed segmentationand further filtered for sphericity (> 0.4), volume (> 000.5 µm^3^) and co-existence of MPP2-, SynCAM1- and vGlut1 staining within the same synapse (2.5 µm distance cut-off to the centre of the next PSD-95 cluster). M

#### Radial intensity profiles

A custom-written ImageJ [46]script (https://github.com/ngimber/RadialProfile3D) was used to calculate 3D radial intensity profiles around PSD-95 centres (segmentation from above). Radial intensity profiles were summarized, 0-1 normalised and averaged twice (per image and per experiment) using Python before plotting with Prism 7 (GraphPad).

#### Nearest neighbour analysis

Nearest neighbour analysis and randomizations were performed in Python using custom-written scripts. Nearest neighbour distances between PSD-95, MPP2 and SynCAM1 cluster were calculated based on the Arivis segmentations. Random controls were generated by randomly distributing spherical objects, representing PSD-95, MPP2 and SynCAM1 clusters within a simplified spherical post-synapse with a diameter of 0.8 µm[19]. Randomised distributions were generated for each image using the object counts and volumes from the corresponding segmentation (10 simulation rounds per synapse, ∼ 40.000 synapses from 50 images). Plotting was done with Prism 7 (GraphPad).

#### Object statistics

Object counts and sizes were obtained from the segmentation above (Arivis). Histograms (bin size = 15 nm) from cluster sizes were plotted in R+.

### Cell culture and transfection

HEK293T and CHL V79 cells were maintained in low-glucose DMEM supplemented with 10% FCS, 1000 U/ml penicillin/streptomycin and 2 mM L-glutamine in a humidified incubator at 37 °C with 5% CO_2_. Transfections were performed using Lipofectamine 2000 (Invitrogen) and desired DNA constructs diluted in Opti-MEM (Gibco).

### DNA constructs

N-terminal FLAG-tagged MPP2 and PSD-95 were cloned as described before [13, 49]. Full length mouse MPP2 (NM_016695) was cloned into pEGFP-C1, to obtain N-terminal EGFP-tagged EGFP-MPP2. Full-length rat SynCAM1 (NM_001012201.1) was synthesised as described before [13]. Clover-PSD-95. Expression constructs for Flag-tagged full-length rat proteins were generated by cloning Arhgef2 (NM_001012079), Ppp3ca (NM_017041) and Farp1 (NM_001107287) into pCMV-Tag 2A. HA-Gnao1 (NM_017327) was constructed by PCR with a forward primer that encodes the HA tag and cloned with NotI and SalI into pCMV-Tag 2A.

GFP-PIPK1 gamma 90 was a gift from Pietro De Camilli (Addgene plasmid # 22299; http://n2t.net/addgene:22299; RRID: Addgene_22299). GABA (A) receptor subunit a1SE was a gift from Tija Jacob & Stephen Moss (Addgene plasmid # 49168; http://n2t.net/addgene:49168; RRID: Addgene_49168). Rat GABA_A_α1 (NM_183326) was cloned into pCMV-2A with NotI and SalI to generate an untagged expression construct.

The bacterial expression construct GST-MPP2-SH3-GK was generated by cloning the fragment encoding amino acids 220-552 (SH3GK) of mouse MPP2 into pGEX-6P-1 (GE Healthcare). GST-PSD-95-SH3GK was generated as described before [25]. A fragment corresponding to amino acids 403-724 of PSD-95 was cloned into pGEX-6P-1.

### Co-immunoprecipitation

HEK293T cells were harvested 18-20 hrs after transfection with a cell scraper and cell lysates obtained with 30 gauge syringe needle strikes in immunoprecipitation buffer (50 mM Tris pH 7,4; 100 mM NaCl; 1 mM EDTA, 1% Triton-X or 0.1% NP-40; supplemented with Complete Mini protease inhibitors, Roche). Cell lysates were cleared by centrifugation (3x 10 min at 20000 *g* at 4°C) and supernatants were incubated with 2 µg αGFP (mouse, 75-131, NeuroMab), αGABA-Aα1 (mouse, 75-136, NeuroMab) or normal mouse IgG, respectively, for 3 hrs on a rotator (10 rpm) at 4°C. Pull-down was performed with 30 µl protein-G-agarose bead slurry (Roche) for 1 hr at 4°C under gentle agitation, followed by three washes with IP buffer and final analysis by western blot.

### Western Blotting

Lysates and beads were boiled in 2x SDS-sample buffer (8% SDS, 40% glycerol, 0,25 M Tris pH 6.8, 20% β-mercaptoethanol) for 5 min, separated on a 10% SDS-PAGE and blotted onto PVDF membranes (Roche). Membranes were blocked with 5% skim milk/PBST. Primary antibodies were diluted in the blocking buffer and applied over night at 4°C, followed by three washes with PBST and 1 hr incubation with appropriate secondary antibodies. HRP-signals were detected using Western Lightning chemiluminescent substrates (Perkin Elmer) with a luminescent image analyser (ImageQuant LAS4000, GE Healthcare).

#### Primary antibodies

αSynCAM (rabbit, PA3-16744, Thermo Scientific), αMPP2 (rabbit, ab97290, Abcam), αPSD-95 (mouse, 75-028, NeuroMab), αGABA_A_ α1 (mouse, 75-136, NeuroMab), αFlag (mouse, F1804, Sigma), αFlag-HRP (mouse, A8592, Sigma), αGFP (chicken, ab13970, Abcam), αGST (mouse, 75-148, NeuroMab).

#### Secondary antibodies

αMouse-HRP (115-035-003, Dianova), αMouse-native-IgG (Veriblot, 131368, Abcam), αRabbit-HRP (111-035-003, Dianova), αRat-HRP (sc-2032, Santa Cruz), αChicken-HRP (ab6753, Abcam).

### Crude synaptosome fraction preparation

For immunoprecipitation using brain lysates, adult Wistar rats were administered isofluorane anaesthesia prior to decapitation and reported under permit T0280/10 (LaGeSO). The brains were removed and rinsed in ice-cold PBS. Brains were immediately frozen and stored at −80°C until use. Brains (∼2 g) were then thawed on ice, minced with a scalpel and homogenized in 20 ml Syn-PER Synaptic Protein Extraction Reagent (Thermo Science) according to the manufacturer’s protocol. For co-immunoprecipitation, the resulting crude synaptosome fraction was then resuspended in IP buffer (50 mM Tris pH 7,4; 100 mM NaCl; 1 mM EDTA, 1% Triton-X; supplemented with Complete Mini protease inhibitors, Roche) and cleared by 3x centrifugation at 20000 *g*. For GST pull-down, the pellet was solubilised in 10 ml 1% Triton-X/PBS.

### GST pull-down

GST-SH3-GK domain constructs of PSD-95 and MPP2 were expressed in *E*.*coli* BL21 DE3 and purified according to the manufacturer’s manual (GST Gene Fusion System, GE Healthcare). 30 µl of glutathione agarose (Pierce) was loaded with GST-SH3-GK proteins (PSD-95 and MPP2) and incubated for 3 hrs with protein lysate from crude synaptosomes. The beads were washed three times with PBS/1% Triton X-100 and eluted from the matrix by incubation with SDS sample buffer for 5 min at 95°C.

### Sample preparation and liquid chromatography-mass spectrometry (LC-MS)

Proteins from two technical replicates were separated by SDS-PAGE (10% Tricine-SDS-PAGE). Coomassie-stained lanes were cut into 12 slices and in-gel protein digestion and ^16^O/^18^O-labelling was performed as described previously [50, 51]. In brief, corresponding samples (PSD-95 and MPP2) were incubated overnight at 37°C with 50 ng trypsin (sequencing grade modified, Promega) in 25 µl of 50 mM ammonium bicarbonate in the presence of heavy water (Campro Scientific GmbH, 97% ^18^O) and regular ^16^O-water, respectively. Isotope-labels were switched between the two replicates. To prevent oxygen back-exchange by residual trypsin activity, samples were heated at 95°C for 20 min. After cooling down, 50 µl of 0.5% trifluoroacetic acid (TFA) in acetonitrile was added to decrease the pH of the sample from pH8 to pH2. Afterwards, corresponding heavy- and light-isotope labelled samples were combined and peptides were dried under vacuum. Peptides were reconstituted in 10 µl of 0.05% TFA, 2% acetonitrile, and 6.5 µl were analysed by a reversed-phase nano liquid chromatography system (Ultimate 3000, Thermo Scientific) connected to an Orbitrap Velos mass spectrometer (Thermo Scientific). Samples were injected and concentrated on a trap column (PepMap100 C18, 3 µm, 100 Å, 75 µm i.d. x 2 cm, Thermo Scientific) equilibrated with 0.05% TFA, 2% acetonitrile in water. After switching the trap column inline, LC separations were performed on a capillary column (Acclaim PepMap100 C18, 2 µm, 100 Å, 75 µm i.d. x 25 cm, Thermo Scientific) at an eluent flow rate of 300 nl/min. Mobile phase A contained 0.1% formic acid in water, and mobile phase B contained 0.1% formic acid in acetonitrile. The column was pre-equilibrated with 3% mobile phase B followed by an increase of 3–50% mobile phase B in 50 min. Mass spectra were acquired in a data-dependent mode using a single MS survey scan (m/z 350–1500) with a resolution of 60,000 in the Orbitrap, and MS/MS scans of the 20 most intense precursor ions in the linear trap quadrupole. The dynamic exclusion time was set to 60 s and automatic gain control was set to 1 × 10^6^ and 5,000 for Orbitrap-MS and LTQ-MS/MS scans, respectively.

### Proteomic data analysis

Identification and quantification of ^16^O/^18^O-labelled samples was performed using the Mascot Distiller Quantitation Toolbox (version 2.7.1.0, Matrix Science). Data were compared to the SwissProt protein database using the taxonomy rattus (August 2017 release with 7996 protein sequences) including the sequences of the employed protein constructs and the sequence of the GST tag. A maximum of two missed cleavages was allowed and the mass tolerance of precursor and sequence ions was set to 15 ppm and 0.35 Da, respectively. Methionine oxidation, acetylation (protein N-terminus), propionamide (Cysteine), and C-terminal ^18^O_1_- and ^18^O_2_-isotope labelling were used as variable modifications. A significance threshold of 0.05 was used based on decoy database searches. For quantification at protein level, a minimum of two quantified peptides was set as a threshold. Protein ratios were calculated from the intensity-weighted average of all corresponding peptide ratios. The protein ratio of GST was used for normalisation of protein ratios. Only proteins that were quantified in both replicates with a standard deviation of <2 were considered. Known contaminants (e.g. keratins) and the bait proteins were removed from the protein output table.

## Acknowledgements

We are grateful for technical assistance from Melanie Fuchs. For mass spectrometry, we would like to acknowledge the assistance of the Core Facility BioSupraMol.

## Supplemental information captions

**S1 Fig: Dual colour dSTORM images of SynCAM1 and PSD-95.**

E18 rat primary hippocampal neurons were fixed at DIV21 and stained for endogenous SynCAM1 (magenta) and PSD-95 (cyan) proteins with Alexa Fluor 647 and Cy3b-coupled secondary antibodies. Protein localizations were filtered according to Thompson accuracy, i.e. all localisations with accuracy below 20 nm were excluded. Box size = 750 nm.

**S2 Fig: Dual colour dSTORM images of MPP2 and PSD-95.**

E18 rat primary hippocampal neurons were fixed at DIV21 and stained for endogenous MPP2 (magenta) and PSD-95 (cyan) proteins with Alexa Fluor 647 and Cy3b-coupled secondary antibodies. Protein localizations were filtered according to Thompson accuracy, i.e. all localisations with accuracy below 20 nm were excluded. Box size = 750 nm.

**S3 Fig: Dual colour dSTORM images of SynCAM1 and MPP2.**

E18 rat primary hippocampal neurons were fixed at DIV21 and stained for endogenous SynCAM1 (magenta) and MPP2 (cyan) proteins with Alexa Fluor 647 and Cy3b-coupled secondary antibodies. Protein localizations were filtered according to Thompson accuracy, i.e. all localisations with accuracy below 20 nm were excluded. Box size = 750 nm.

**S4 Fig: Nearest Neighbour Analysis of SynCAM1 and MPP2 protein cluster distances derived from SIM images.** NN distances from SynCAM1 to the nearest MPP2 cluster were calculated from the cluster centres (upper panel, grey bars) and cluster surfaces (lower panel). Dashed lines represent the upper and lower envelopes of complete spatial randomness (CSR). CSR was calculated by randomly distributing MPP2 within the volume and SynCAM1 on the surface of spheres of 0.8 µm diameter as indicated by the grey dotted line (mean ± SEM, 95% confidence interval, 10 simulations per synapse, N = 3 independent experiments, ∼40.000 synapses from 50 images).

**S5 Fig: Validation of novel interaction partners by Co-IP**

a) HA-tagged Gnao1 was overexpressed together with EGFP-tagged MPP2 in HEK293T cells. Upon pull-down with Ms αGFP antibody or normal Ms IgG, Gano1 co-purification and GFP pull-down control were detected by Western blot with αHA and αGFP antibodies.

b) Co-IP of FLAG-tagged Arhgef2 together with EGFP-MPP2 after pull-down with αGFP antibody or IgG control, as detected by Western blot using αFLAG and αGFP antibodies.

c) FLAG-tagged Farp1 was overexpressed together with EGFP-tagged MPP2 and co-purifies with αGFP pull-down, as opposed to normal IgG as negative control. Co-IP was detected by Western blot probing with αFLAG and αGFP antibodies.

d) Co-purification of FLAG-tagged Ppp3ca (Calcineurin subunit) overexpressed together with EGFP-MPP2 after αGFP pull-down or normal Ms IgG as negative control, detected by Western blot with αFLAG and αGFP antibodies.

e) EGFP-tagged Pip5k1c was co-expressed with FLAG-tagged MPP2 in HEK293T cells. EGFP-Pip5k1c was precipitated with αGFP antibody or normal IgG as negative control and analysed by Western blot with αFLAG and αGFP antibodies. An additional IgG control lane is marked with an asterisk.

**S6 Fig: Full-length blots for Fig 4**.

**S7 Fig: Full-length blots for S5 Fig**.

**S1 Data: Underlying source data for Fig 3c**.

